# Cigarette smoke exposed airway epithelial cell-derived EVs promote pro-inflammatory macrophage activation in alpha-1 antitrypsin deficiency

**DOI:** 10.1101/2022.07.07.499205

**Authors:** Nazli Khodayari, Regina Oshins, Borna Mehrad, Jorge E. Lascano, Xiao Qiang, Jesse R. West, L. Shannon Holliday, Jungnam Lee, Gayle Wiesemann, Soroush Eydgahi, Mark Brantly

**Author notes:** **Corresponding author:** Nazli Khodayari, Assistant Professor, University of Florida, College of Medicine, 1600 SW Archer Rd Rm M453A, Gainesville, FL 32610 USA. **Funding:** Alpha-1 Foundation (F007320).

## Abstract

**Background:** Alpha-1 antitrypsin deficiency (AATD) is a genetic disorder most commonly secondary to a single mutation in the SERPINA1 gene (PI*Z) that causes misfolding and accumulation of alpha-1 antitrypsin (AAT) in hepatocytes and mononuclear phagocytes which reduces plasma AAT and creates a toxic gain of function. This toxic gain of function promotes a pro-inflammatory phenotype in macrophages that contributes to lung inflammation and early-onset COPD, especially in individuals who smoke cigarettes. The aim of this study is to determine the role of cigarette exposed AATD macrophages and bronchial epithelial cells in AATD-mediated lung inflammation.

**Methods:** Peripheral blood mononuclear cells from AATD and healthy individuals were differentiated into alveolar-like macrophages and exposed to air or cigarette smoke while in culture. Macrophage endoplasmic reticulum stress was quantified and secreted cytokines were measured using qPCR and cytokine ELISAs. To determine whether there is “cross talk” between epithelial cells and macrophages, macrophages were exposed to extracellular vesicles released by airway epithelial cells exposed to cigarette smoke and their inflammatory response was determined.

**Results:** AATD macrophages spontaneously produce several-fold more pro-inflammatory cytokines as compared to normal macrophages. AATD macrophages have an enhanced inflammatory response when exposed to cigarette smoke-induced extracellular vesicles (EVs) released from airway epithelial cells. Cigarette smoke-induced EVs induce expression of GM-CSF and IL-8 in AATD macrophages but have no effect on normal macrophages. Release of AAT polymers, potent neutrophil chemo attractants, were also increased from AATD macrophages after exposure to cigarette smoke-induced EVs.

**Conclusions:** The expression of mutated AAT confers an inflammatory phenotype in AATD macrophages which disposes them to an exaggerated inflammatory response to cigarette smoke-induced EVs, and thus could contribute to progressive lung inflammation and damage in AATD individuals.

## Background

Alpha-1 antitrypsin deficiency (AATD) is a genetic illness caused by a single nucleotide mutation in the SERPINA1 gene (1). AATD is the most common genetic risk factor for chronic obstructive pulmonary disease (COPD) and individuals who smoke die 20 years before non-smokers (2). COPD is an irreversible inflammatory airway disease characterized by airflow obstruction and emphysema, usually caused by smoking (3, 4). Alpha-1 antitrypsin (AAT) is a protease inhibitor synthesized mainly by hepatocytes and, to a lesser extent, by monocytic phagocytes (1). The normal variant of AAT (M variant, MAAT) is secreted into the circulation, where its primary function is to protect different tissues against a wide range of proteases, such as neutrophil elastase (NE). The mutant Z variant of AAT (ZAAT), characterized by a single amino acid substitution of lysine for glutamic acid at position 342, is prone to misfolding, aggregation, and accumulation within the endoplasmic reticulum (ER) of hepatocytes and monocytic phagocytes. Resulting low levels of circulating AAT cause pulmonary inflammation due to uncontrolled proteolytic activity of proteases and excessive degradation of lung parenchyma, particularly in response to cigarette smoke exposure (2). AATD is the cause of 1–2% of COPD cases (5).

The population of lung macrophages is expanded in patients with COPD as compared to healthy subjects, and this expansion correlates with the severity of lung disease (6). While alveolar macrophages have an immunosuppressive phenotype in healthy individuals, recent evidence indicates lung macrophages display a pro-inflammatory phenotype in COPD patients (7). The ligands of several pattern recognition receptors, inhibition of IL-10 receptor signaling, and activation of NF-κB signaling have been shown to trigger the pro-inflammatory state in alveolar macrophages (8). Pro-inflammatory macrophages display impaired phagocytic activity and secretion of pro-inflammatory cytokines. In COPD, pro-inflammatory macrophages promote disease progression by releasing high levels of pro-inflammatory cytokines, including IL-8, driving recruitment of neutrophils and monocytes to the lungs (9). Furthermore, increased numbers of lung neutrophils result in a profound proteolytic burden in the lung of AATD individuals (10).

Both alveolar and monocyte-derived macrophages express and secrete AAT. Monocyte differentiation to macrophages has been shown to increase the expression level of AAT up to 3-fold (11). In AATD, accumulation of misfolded ZAAT in monocytes and macrophages causes the unfolded protein response and activates NF-kb pathways and expression of pro-inflammatory cytokines (11). The ZAAT polymers within the lungs are also a potent pro-inflammatory chemoattractant agent for human neutrophils. It is therefore likely that ZAAT polymers contribute to the increased inflammation in the lung of AATD individuals (12).

Extracellular vesicles (EVs), which include exosomes and microvesicles, are 30–150 nm diameter membrane-bound vesicles released by all types of cells and have roles in intercellular signaling and modulation of cellular homeostasis (13). EVs enriched with pro-inflammatory cargo contribute to lung inflammation, suggesting they play a critical role in the inflammation state of pathological conditions, including COPD (14). It has been shown that during the progression of COPD, a large number of EVs can be found in the sputum, plasma and bronchial lavage fluid of patients, regulating immune cells by mediating intercellular communication (15). Exposure to cigarette smoke and air pollutants has been shown to affect the number and cargo of EVs released by different cell populations in the lungs and participates in development of lung diseases (15). Airway epithelial cells, the major cell population exposed to cigarette smoke (16), release EVs with distinct cargo that modulates the activation of macrophages within the lungs during the inflammatory state of COPD (13, 16). Whether AATD associated inflammation is modulated by EVs remains to be determined, as does the contribution of EVs to immune responses in AATD-mediated COPD.

Expression of misfolded ZAAT and subsequent activation of the ER stress pathways in monocytes from AATD individuals have been investigated previously (17). Furthermore, we have previously shown that peripheral blood monocyte-derived macrophages from AATD individuals have impaired efferocytosis (5) and dysregulated proteolytic activity (2). Taken together, these data suggest an altered phenotype of AATD macrophages could be involved in the pathogenesis and severity of AATD-mediated COPD. We therefore hypothesized that enhanced inflammatory phenotype of AATD macrophages and dysregulated response to cigarette smoke and cigarette smoke-induced EVs contribute to the development of lung inflammation in AATD individuals.

## Materials and methods

### Subjects

Peripheral blood mononuclear cells expressing M or Z variants of AAT were isolated from blood obtained from outpatient volunteers (Table 1) after informed consent (University of Florida Institutional Review Board protocol # 2015-01051). All individuals were healthy at the time of blood collection.

**Table 1.** Characterization of normal and AATD subjects.

| Characteristic | Normal Individual<br>(n=6) | AATD Individual<br>(n=6) |
| --- | --- | --- |
| Genotype | MM | ZZ |
| Plasma AAT (μM) | 25.6 (20.4 - 35.0) | 3.7 (2.9 - 5.8) |
| Age, Year | 35.8 (22-48) | 45 (36-62) |
| Male sex | 3 | 2 |
| FEV<sub>1</sub>% | 98.3 (71-116) | 73.5 (52-96.6) |
| Current smoker | None | None |
| Lung Disease | None | Emphysema (5/6) |

### Cell culture

Macrophages were generated from peripheral blood mononuclear cells as previously described (2). Peripheral blood mononuclear cells were isolated from the blood of outpatient volunteers (University of Florida, IRB # 2015-01051) using Ficoll gradient centrifugation. Approximately 400 mL of whole blood was collected and centrifuged at 2600 × g for 10 minutes. Plasma was removed and white blood cells were combined with PBS containing 2 mM EDTA at a ratio of 1:1. The cells were overlaid onto Ficoll-Paque Plus (GE Healthcare, Chicago, IL) at a ratio of 2.8:1 and centrifuged at 400 × g for 30 minutes to remove red blood cell contamination. The white blood cells were collected and washed twice with PBS and plated in serum-free RPMI. After 2 hours, unattached cells were discarded and adherent monocytes were differentiated into macrophages by culturing for 7 days in RPMI with 10% FBS, 20 Units/mL penicillin, 20 μg/mL streptomycin, 250 ng/mL Amphotericin B, 1 ng/mL GM-CSF and 10 ng/mL M-CSF. To characterize differentiated macrophages, they were compared to macrophages polarized with GM-CSF or M-CSF only as we have previously reported (2).

Primary human small airway epithelial cells were purchased from ATCC (Manassas, VA). Small airway epithelial cells were maintained in airway epithelial cell basal medium (ATCC, PCS-300-030) supplemented with the small airway epithelial cell growth kit (ATCC, PCS-301-040) at 37°C in an atmosphere of 95% air supplemented with 5% CO_2_ (18).

### Cigarette smoke and EV exposure

Macrophages and early passage small airway epithelial cells were exposed to one exposure unit (smoke exposure from three whole cigarettes over 15 minutes followed by 45 minutes of incubation in 5% CO_2_ and 95% ambient air) of 3R4F research cigarettes (Kentucky Tobacco Research and Development Center) or air, as we have previously published (19). Briefly, the inlet of a modular incubator chamber (Billups-Rothenberg, CA) was connected to a cigarette holder in a fume hood. To mimic in-vivo alveolar lung fluid, monolayers of cultured cells on the bottom of a 100-mm culture dish were covered by 8 mL medium. The culture dishes in the chamber were then exposed to cigarette smoke. The cells in a parallel chamber were treated under identical conditions, but without cigarette smoke and used as non-smoked controls. Forty-five minutes after exposure, the media was changed to fresh RPMI with 10% EV-free FBS or supplemented airway epithelial cell basal media. The morphology of the cells was monitored by light microscopy (Supplementary Figure 1A, and Figure 2A). Cells and media were collected after 8 and 24 hours for further analysis.

For incubation with EVs, normal and AATD macrophages were plated at a cell concentration of 200,000 per well in 12-well plates. Isolated EVs (1 × 10^7^ control or smoked airway epithelial cell derived EVs) were added to each well and the plate was incubated for 8 or 24 hours at 37°C. Normal and AATD macrophages incubated with an equivalent volume of PBS have been used as negative control.

### EV isolation and characterization

EV isolation and preparation from airway epithelial cell and macrophage conditioned media was performed as we previously described (20). Briefly, conditioned media were collected 8 or 24 hours post cigarette smoke exposure and centrifuged at 1,000 x g for 10 minutes to remove dead cells and cellular debris. The supernatant was then filtered with a 0.22-*μ*m Nalgene filter (Thermo Scientific). The resulting supernatant was centrifuged at 10,000 x g for 30 minutes to remove any remaining debris and then ultra-centrifuged at 118,000 x g for 70 minutes at 4°C with a fixed-angle rotor (Ti-70, Beckman Coulter, Brea, CA). The pellet was washed with sterile 1X PBS and subjected to another cycle of ultracentrifugation at 118,000 x g for 70 minutes at 4°C. The supernatant was discarded and the pelleted EVs were carefully reconstituted in sterile 1X PBS or lysed in RIPA buffer (ThermoFisher Scientific, Carlsbad, CA). EV concentration and characterization were determined using Nano Sight-based EV technology. EV morphology was characterized by imaging on a HITACHI 7600 transmission electron microscope equipped with AMTV600 camera at the University of Florida Transmission Electron Microscopy core (20).

### EV labeling and uptake assay

For uptake analysis, purified EVs from conditioned media were incubated with Fast DiO membrane dye (Invitrogen, Carlsbad, CA) at a final concentration of 2 μg/mL for 1 hour at room temperature. The purification process of washing and ultracentrifugation was repeated twice, and the labeled EV pellet was resuspended in PBS. For microscopic analysis, normal macrophages and AATD macrophages were incubated with DiO-labeled EVs for 1 hour at 37°C. After incubation, cells were treated with trypsin followed by PBS to remove unbound labeled EVs and subsequently imaged with a Keyence fully motorized BZ-X800 microscope (KEYENCE America, Chicago, IL).

### Cytokine measurement assay

We identified priority cytokine candidates based on known macrophage cytokine profiles and previously published literature, then selected from assays available at Myriad-RBM, which were primarily multiplexes (Luminex, Myriad-RBM Inc., Austin TX). Conditioned media was run on the MilliPLEX Human High Sensitivity T Cell Magnetic Bead Panel Luminex kit for measurement of 21 unique cytokines per plex according to manufacturer’s instructions (EMD Millipore, St. Louis, MO).

### Quantitative real-time PCR

Total RNA was extracted from both normal and AATD macrophages, and the first complementary strand was generated. The expression level of ATF4, XBP1 and CHOP gene products was analyzed by quantitative real-time PCR using an Applied Biosystems 7500 fast real-time PCR system (Life Technologies, Carlsbad, CA) and TaqMan universal PCR master mix from Roche Applied Science. Pairs of genes were analyzed simultaneously, and h18S ribosomal RNA used as an endogenous control. Results are presented as relative quantification determined by the 2−ΔΔCt equation as previously reported (21).

### Western blot analysis

Macrophages and cultured media were lysed in PBS or RIPA buffer containing protease inhibitor cocktail and were separated by native non-denaturing or denaturing 10% SDS-PAGE and transferred to a nitrocellulose membrane. Membranes were blotted with antibody specific to AAT (DAKO, Carpinteria, CA), Ik*β*, p-Ikk*β* and *β*-actin (Cell Signaling, Danvers, MA), Calnexin, CD63, TSG101, TNF-*α*, IL-1*β* (Proteintech, Chicago, IL), and GAPDH (Santa Cruz Biotechnology, Dallas, TX, USA).

### Immunofluorescence assay

To examine AAT distribution within cultured macrophages, they were differentiated on glass slides and fixed in 4% paraformaldehyde for 15 minutes followed by permeabilization in PBS containing 0.01% Triton X-100. The permeabilized macrophages were then incubated with antibody against AAT (Dako, Carpinteria, CA) and were immunostained with Alexa Fluor488 secondary antibody (Abcam, Cambridge). The immunostained cells were mounted on glass coverslips using VECTASHIELD mounting media with DAPI and examined using a fluorescence microscope (BZ-X800, KEYENCE America, Chicago, IL) (5).

### Statistical Analysis

All results are presented as the mean ± SE. Statistical analysis was performed using the two-tailed Student’s t test (GraphPad Prism 9: GraphPad Software, San Diego, CA). *p* values less than 0.05 were considered statistically significant.

## Results

### Characterization of AATD macrophages

After 7 days of culture, no significant differences in morphology between normal and AATD macrophages were detected (Figure 1A). We found no difference in mRNA expression levels of AAT in normal and AATD macrophages by qPCR analysis (Figure 1B). However, macrophages from AATD individuals had higher intracellular protein levels of AAT as measured by western blot analysis (Figure 1C) and immunofluorescent staining (Figure 1D and E) compared to cells from healthy donors. In addition, using the Luminex cytokine assay, we observed higher concentrations of CCL2, CCL3, CCL4, TNF-*α* and IL-6 in the conditioned media from AATD macrophages (n=6) as compared to normal macrophages under resting conditions, p < 0.05-0.0005 (Figure 1F).

**Figure 1.**
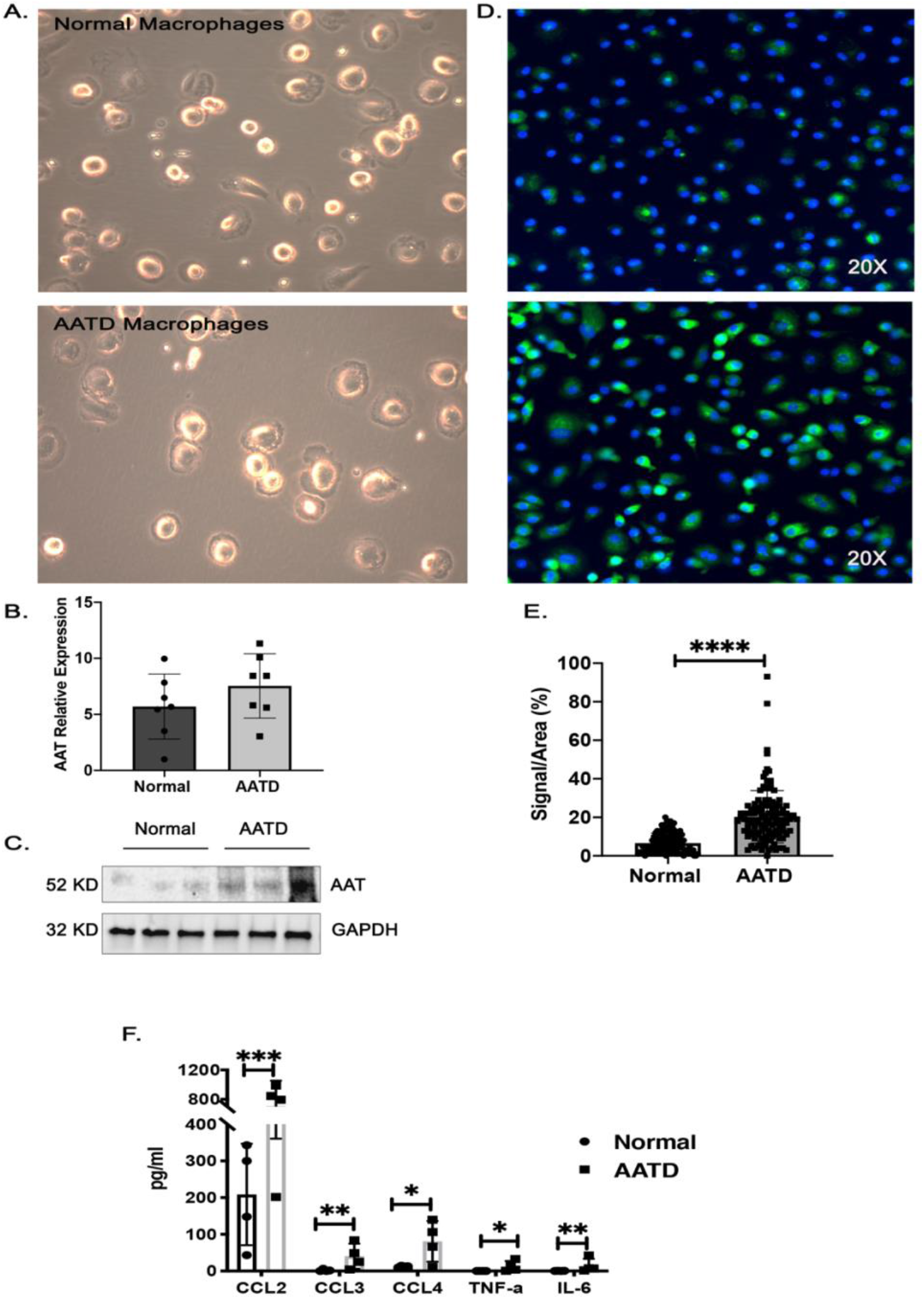
Characterization of normal and AATD macrophages. (A) The morphology of monocyte-derived macrophages from normal and AATD individuals was visualized by light microscopy. (B) The relative expression of AAT mRNA from normal and AATD macrophages. (C) The intracellular protein levels of AAT in representative macrophages. (D) Immunofluorescent images of normal and AATD macrophages at 20X magnification demonstrate intracellular AAT levels using FITC-labeled AAT antibody. (E) Quantification of the intracellular levels of AAT protein. (F) Comparison of the cytokine levels in the culture media of normal and AATD macrophages, *: p<0.05, **: p<0.005, ***: p<0.0005, ****: p<0.00005.

### Changes in the macrophage’s cytokine secretion profile in response to cigarette smoke

To characterize the impact of cigarette smoke exposure on cytokine production by normal and AATD macrophages, we measured the levels of secreted cytokines in the media of cultured normal and AATD macrophages without treatment and 24 hours post cigarette smoke exposure. First, cytotoxicity and viability were measured using LDH and MTT assays. LDH assay indicated no cytotoxicity (Supplementary Figure 2B) due to cigarette smoke exposure. In contrast MTT assay showed significant decrease in the viability of AATD macrophages after exposure to cigarette smoke while normal macrophages showed very mild changes after exposure to cigarette smoke (Supplementary Figure 2C). Furthermore, we observed that cigarette smoke has a mild inhibitory effect on the cytokine production by both normal and AATD macrophages as previously reported (22). Although cigarette smoke dysregulated the cytokine production in both normal and AATD macrophages, we were unable to detect any significant differences in response to cigarette smoke between normal and AATD macrophages (Figure 2A and B). We also determined the levels of EV-associated cytokines produced by normal and AATD macrophages in response to cigarette smoke. To do this, we first analyzed the concentration of EVs released by normal and AATD macrophages in response to cigarette smoke and found that there are no significant differences in the release of EVs from normal and AATD macrophages after cigarette smoke exposure (Supplementary Figure 1C). Consistent with the levels of free cytokines produced by macrophages in response to cigarette smoke, we found cigarette smoke exposure equally effects and dysregulates the levels of EV-associated cytokines released from normal and AATD macrophages (Figure 2C and D).

**Figure 2.**
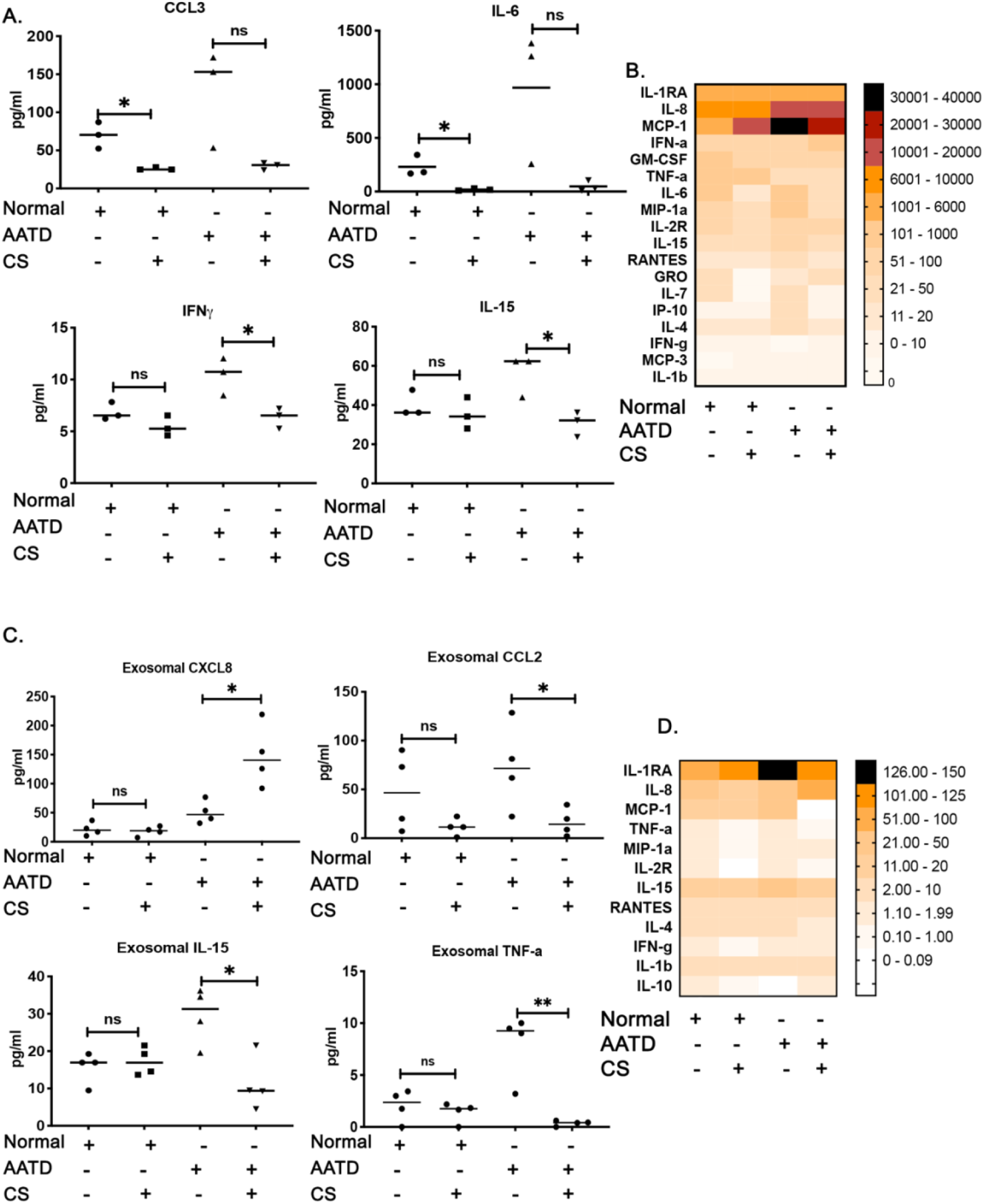
Changes in the macrophage’s cytokine profile in response to cigarette smoke. (A) The cytokine levels in the culture media of normal and AATD macrophages exposed to cigarette smoke (CS) or air. (B) Heatmap presentation of the cytokines. (C) The levels of cytokines associated with EVs in the cultured media of normal and AATD macrophages exposed to CS or air. (D) Heatmap presentation of the EV-associated cytokines. *: p<0.05, **: p<0.005.

### Induction of ER stress and the NF-κB inflammatory pathway in AATD macrophages in response to cigarette smoke

In smokers, different lung cell populations develop ER stress in response to cigarette smoke exposure (14, 23). To examine the impact of cigarette smoke exposure on ER homeostasis in normal and AATD macrophages, we examined expression levels of ER stress marker genes XBP-1, CHOP, ATF4, and BiP in normal and AATD macrophages before and after cigarette smoke exposure. AATD macrophages have higher basal expression of ER stress associated genes due to AAT accumulation and our experiments indicated that only AATD macrophages showed significant induction of ER stress related genes (p<0.05) in response to cigarette smoke exposure (Figure 3A). We next examined the response of the NF-κB inflammatory pathway in normal and AATD macrophages under control conditions and in response to cigarette smoke. Western blot analysis revealed the activation of the NF-κB pathway in AATD macrophages in response to cigarette smoke exposure. Significantly higher levels of phosphorylated-Ikkβ were measured after cigarette smoke exposure (p<0.05) (Figures 3B and C).

**Figure 3.**
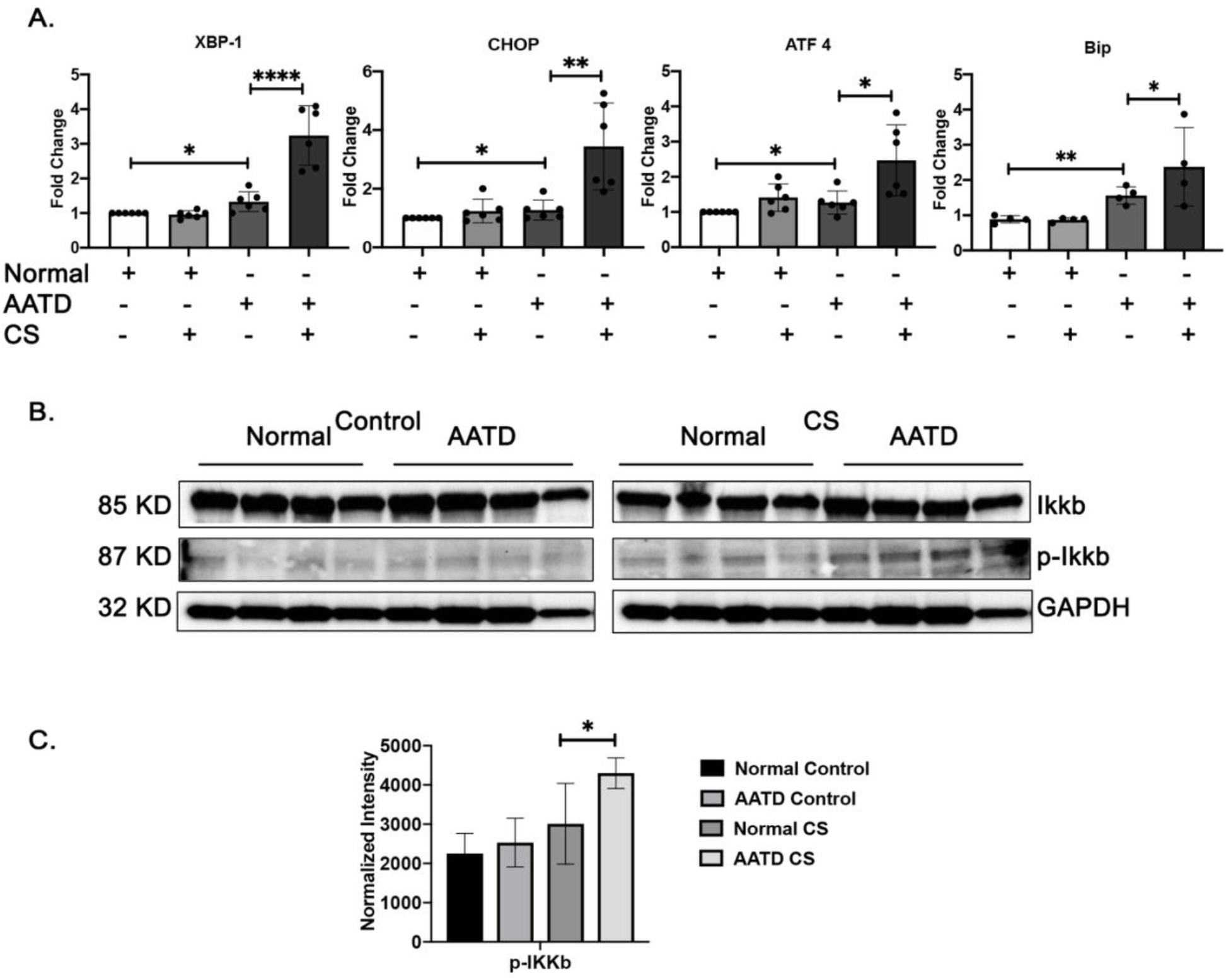
Induction of ER stress and NF-κB inflammatory pathway in AATD macrophages in response to cigarette smoke. (A) Comparison of the mRNA expression levels of XBP-1, CHOP, ATF 4, and BiP in normal and AATD macrophages exposed to air or CS. (B) Western blot analysis of 4 different normal and AATD macrophages exposed to air or CS showing activation of NF-κB pathway in AATD macrophages after CS exposure. (C) Bar graph showing the results of quantification and normalization of band intensities, *: p<0.05, **: p<0.005, ****: p<0.00005.

### Characterization of small airway epithelial cell-derived EVs

To determine whether EVs were secreted by airway epithelial cells and whether there were differences in the size or number from control versus cigarette smoke exposed cells, we assessed size distribution and concentration of EVs in the culture media via Nanoparticle Tracking Analysis (NTA) which demonstrated enrichment of EVs between 50 nm and 200 nm. Airway epithelial cells exposed to cigarette smoke secreted a significantly greater number of EVs per milliliter of media compared with control (Figure 4A). Further evaluation using transmission electron microscopy revealed double membrane cup-shaped vesicles within the expected range of size from 50 nm to 200 nm (black arrows) (Figure 4B). Western blot analysis confirmed abundant CD63, TSG101 and actin in EV fractions and absence of ER markers such as Calnexin. Western blot analysis also indicated that EVs derived from cigarette smoke exposed airway epithelial cells contain more membrane form and soluble TNF-α, as well as pro and active forms of IL-1β compared to control EVs (Figure 4C). We also investigated the levels of EV associated cytokines released from control and cigarette smoke exposed airway epithelial cells. A MILLIPLEX cytokine assay showed few detectable cytokines associated with the EVs. Among those detectable cytokines, the level of EV associated IL-1Ra was completely suppressed in EVs derived from cigarette smoke-induced airway epithelial cells compared to control airway epithelial cells (Figure 4D).

**Figure 4.**
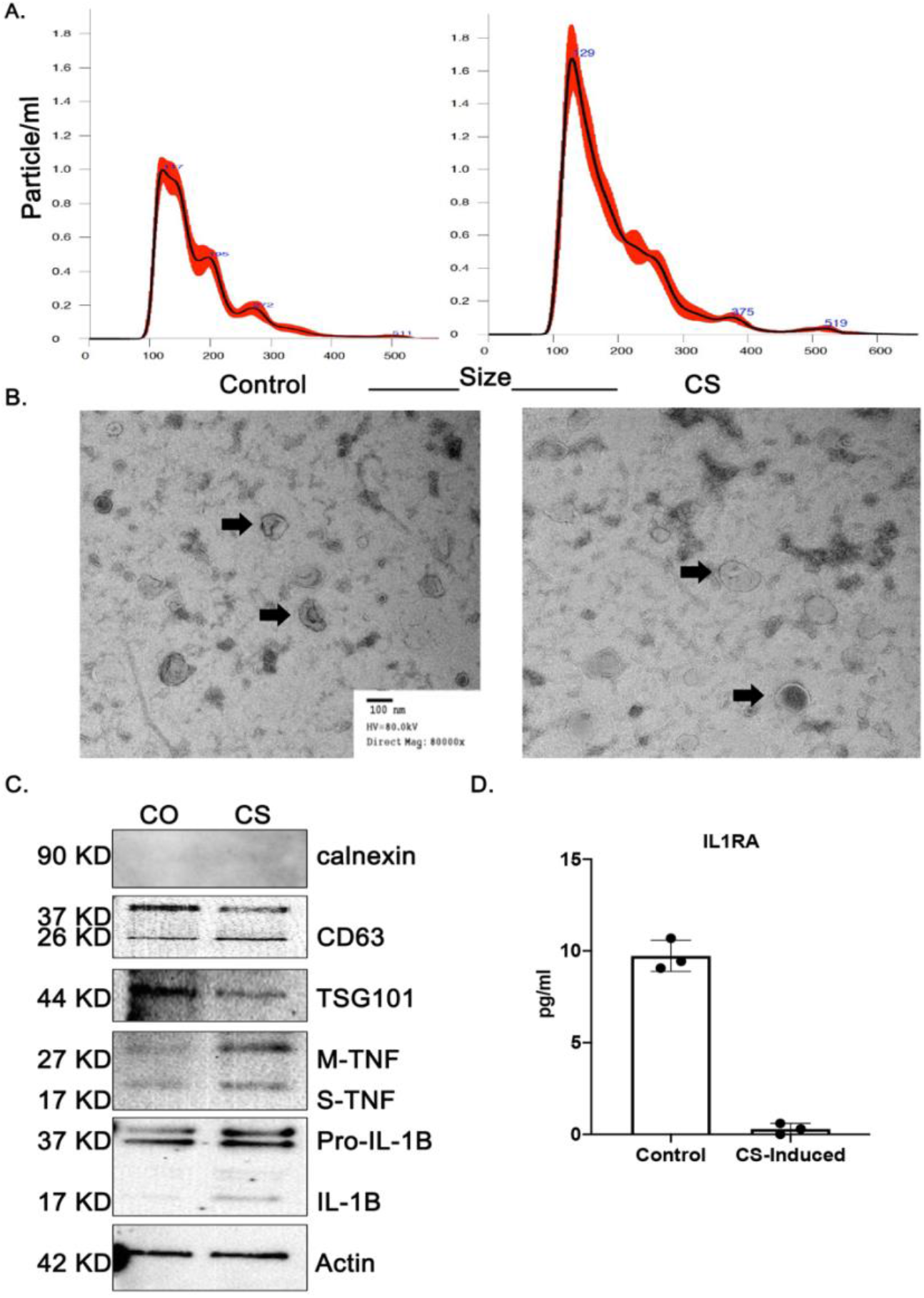
Characterization of airway epithelial cell-derived EVs. (A) Nanoparticle tracking analysis of size and concentration for EVs isolated from conditioned media of control and cigarette smoke-induced EVs. The bold red curves plot the size distribution of EVs. (B) Morphological characterization of EVs by transmission electron microscopy (black arrows). (C) Western blotting experiment of the purified EVs showing Calnexin, CD63, TSG101, membrane bound and soluble TNF-*α* and pro and active IL-1*β*. (D) The levels of cytokines associated with EVs released by small airway epithelial cells.

### AATD macrophage activation toward a pro-inflammatory phenotype and increased secretion levels of ZAAT polymers in response to cigarette smoke-induced EVs

To determine the uptake efficiency of EVs released from airway epithelial cells by normal and AATD macrophages, DiO labeled EVs were incubated with macrophages for 1 hour. Both normal and AATD macrophages had the same rate of intake of EVs (Supplementary Figure 1B). We then investigated secretion levels of several cytokines and chemokines of normal and AATD macrophages in response to exposure to control and cigarette smoke-induced EVs from airway epithelial cells for 8 and 24 hours. It has been previously shown that the NF-κB pathway is an inducible transcriptional activator of IL-8 and GM-CSF genes (24-27). Therefore, we focused on secretion levels of these cytokines and found AATD macrophages had significantly elevated secretion levels of IL-8 and GM-CSF in response to cigarette smoke-induced EVs at both 8 and 24 hours (p<0.005) while there were no significant changes in the cytokine profiles of normal macrophages (Figure 5A). Next, to confirm the role of the NF-κB pathway in response to cigarette smoke-induced EVs in AATD macrophages, we inhibited the NF-κB pathway in AATD mascrophages using 3 μM TPCA-1 (Abcam, Cambridge, UK), a direct inhibitor of the NF-κB pathway (Supplementary Figure 3). After normalizing the cell seeding concentration, we observed that TPCA-1 inhibited overexpression of IL-8 and GM-CSF in response to cigarette smoke-induced EVs in AATD macrophages (Figure 5B). In addition to IL-8 and GM-CSF, we detected significant increases in secretion levels of IL-1Ra, MIP-1a and MIP-1b from AATD macrophages compared to normal macrophages in response to 24 hours of exposure to cigarette smoke induced EVs, which is in line with cytokine profiles of patients with cigarette smoke-induced COPD (28) (Figure 5C). In addition to the above cytokines, AAT polymers have been identified as a chemoattractant factor secreted from macrophages in response to cigarette smoke exposure (29). Thus, we compared the release of AAT polymers from normal and AATD macrophages in response to cigarette smoke-induced EVs. Non-denaturing western blot analysis revealed higher amounts of AAT polymers released by AATD macrophages after exposure to cigarette smoke-induced EVs (p < 0.05) compared to normal macrophages (Figures 5D and E). Total AAT levels in the culture media of AATD macrophages incubated with cigarette smoke-induced EVs were also significantly higher compared with the culture media of AATD macrophages incubated with control EVs (Figure 5F) as measured by ELISA (p < 0.0005). To determine the effect of control and cigarette smoke-induced EVs from airway epithelial cells on the ER status of normal and AATD macrophages, we analyzed the protein expression levels of ER stress markers in macrophages. Our western blot analysis revealed that treatment with cigarette smoke-induced EVs increases the protein levels of ER stress markers in normal macrophages, while AATD macrophages presented elevated levels of ER stress markers as compared to normal macrophages prior to treatment with cigarette smoke-induced EVs (Figure 5G).

**Figure 5.**
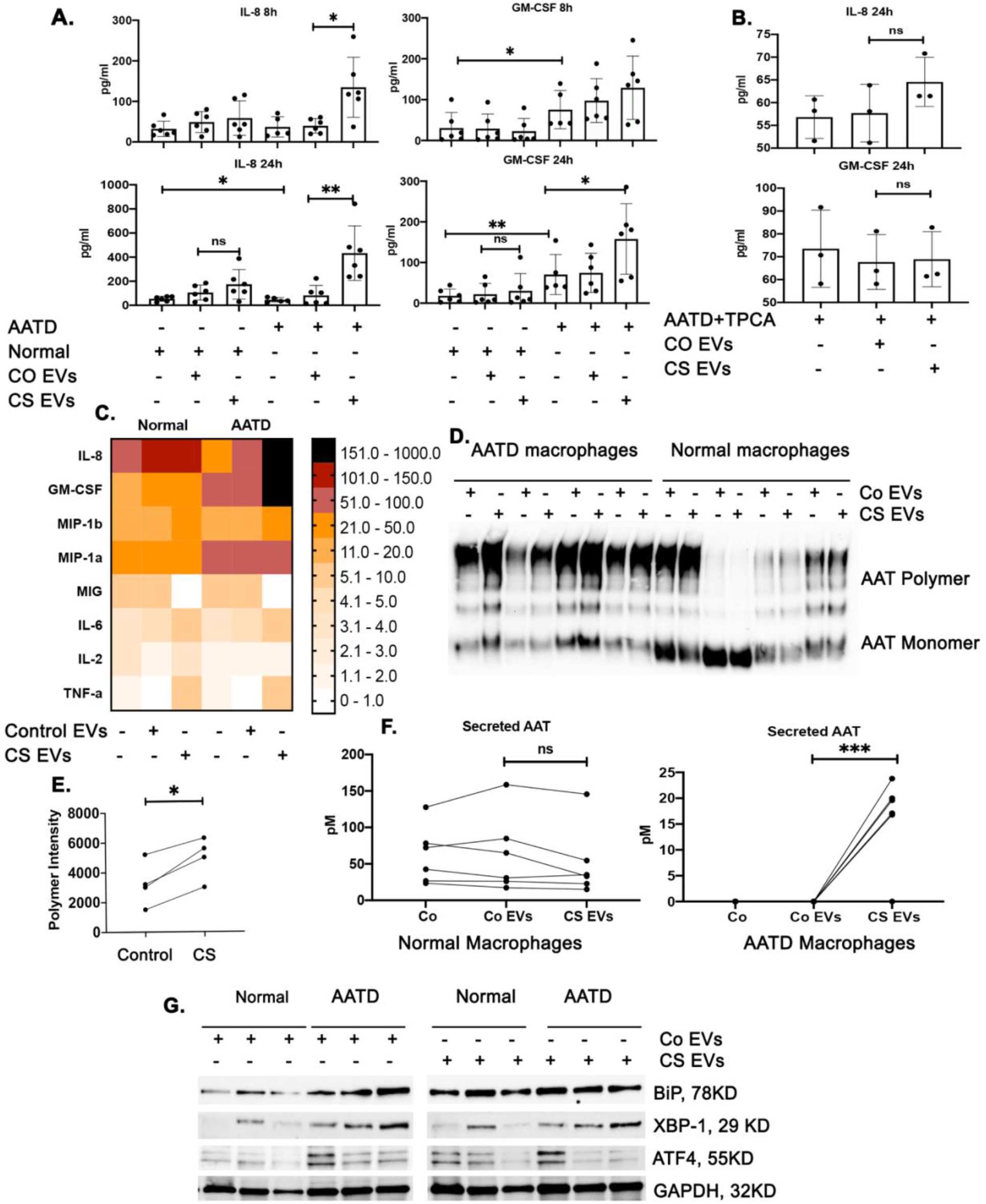
Increased secretion levels of pro-inflammatory cytokines and ZAAT polymers by AATD macrophages in response to cigarette smoke-induced EVs. (A) The levels of secreted cytokines in the conditioned media of normal and AATD macrophages incubated with control or cigarette smoke-induced EVs. (B) NF-κB pathway was inhibited in AATD macrophages using TPCA-1 and the secretion levels of IL-8 and GM-CSF in response to cigarette smoke-induced EVs were measured. (C) The levels of secreted cytokines presented in a heatmap graph. (D) The levels of polymeric AAT in the conditioned media from normal and AATD macrophages incubated with control or cigarette smoke-induced EVs. (E) Signal intensity quantification of polymers in AATD macrophage conditioned media. (F) Total AAT levels were also measured in the conditioned media of normal and AATD macrophages. (G) The protein expression levels of ER markers were measured by western blot analysis. *: p<0.05, **: p<0.005, ***: p<0.0005.

## Discussion

AATD, the only well-established genetic risk factor for COPD, has been thought to solely be the result of low levels of circulating AAT (47); a normal concentration of circulating AAT is 20– 53 μM compared to 3 to 7 μM in AATD individuals (5). The disruption of pulmonary homoeostasis in AATD individuals mostly occurs at an early age and is mainly associated with exposure to cigarette smoke. Two previously reported mechanisms include: low levels of circulating AAT allowing uncontrolled proteolytic activity of neutrophil-derived proteases and tissue destruction; and the presence of misfolded ZAAT in the lung that causes an influx of neutrophils into the lungs of AATD individuals (30). However, the role of macrophage toxic gain of function caused by accumulation of misfolded AAT in the pathophysiology of the disease is still not well established.

Lung macrophages are specialized to create controlled and appropriate immune responses to environmental exposures such as cigarette smoke (31). To maintain pulmonary homoeostasis, it is critical that macrophages sustain an immunotolerant state whilst also being able to rapidly induce effective inflammatory responses (32). We demonstrate that AATD macrophages exhibit a markedly distinct cytokine expression that can contribute to macrophage activation toward a pro-inflammatory phenotype. Moreover, we demonstrate that cigarette smoke disturbs the immune response of AATD macrophages via both direct and indirect effects. We observed that cigarette smoke directly impairs the production of cytokines in macrophages. The pro-inflammatory phenotype of AATD macrophages is also enhanced in response to cigarette smoke-induced EVs released by airway epithelial cells. which is an indirect effect of cigarette smoke on AATD macrophages. We show that elevated levels of ZAAT polymers released from AATD macrophages in response to cigarette smoke-induced EVs may exacerbate lung disease in AATD individuals (30, 33).

In this study, we employed cultured monocyte-derived macrophages (34) and established that AATD macrophages are associated with enhanced secretion levels of the pro-inflammatory cytokines CCL2, CCL3, CCL4, TNF-α and IL-6 compared to normal macrophages under controlled basal conditions. Our results concur with earlier reports in which the levels of CCL2, CCL3 and CCL4 were increased in COPD patients compared to healthy controls and are associated with COPD development (35). Importantly CCL2, which has strong neutrophil chemoattractant activity, is known to have roles in different aspects of the lung inflammatory process, such as tissue remodeling during acute inflammation (36). This may suggest that AATD macrophages are a source of pro-inflammatory cytokines in the lung that fail to resolve inflammation and may contribute to detrimental remodeling in the AATD lung. Lung parenchymal destruction is a key feature of COPD and is a direct result of activated alveolar macrophages (37). Lung tissue destruction results in the loss of lung elasticity and causes airflow obstruction, impairment of defense mechanisms and destruction of alveolar support (38). Our results corroborate these findings, suggesting that the pro-inflammatory phenotype of AATD macrophages might have a role in the increased susceptibility to the development of early age of onset COPD in AATD individuals.

Cigarette smoke is a major risk factor for the development of COPD, inducing persistent lung inflammation. Several studies have indicated that cigarette smoke impairs macrophage function and their secretion of inflammatory cytokines (36). Cigarette smoke is known to induce ER stress and the unfolded protein response (39). ER stress contributes to the pro-inflammatory response in macrophages through the NF-κB signaling pathway (40). Here, we have demonstrated that AATD macrophages have elevated expression levels of ER stress markers at basal conditions compared to normal macrophages. This finding is consistent with previous reports showing accumulation of ZAAT within the ER of AATD macrophages (41). Furthermore, we show that cigarette smoke increases ER stress and significantly affects the viability of AATD macrophages compared to normal macrophages. Consistent with previous studies, we also observed activation of NF-κB in AATD macrophages in response to cigarette smoke. Several studies have suggested that cigarette smoke exposure attenuates cytokine production by alveolar macrophages (42). This effect may be due to the presence of oxygen free radicals and LPS in cigarette smoke damaging signaling pathways upstream of cytokines production (42). In this regard, we also observed that cigarette smoke disturbs cytokine production in both normal and AATD macrophages. However, higher levels of pulmonary neutrophils and macrophages have been observed in mouse models of AATD exposed to cigarette smoke, inducing an inflammatory environment in the lungs (43) which points to the role of AATD macrophages during the developmental course of COPD.

In addition to macrophages, cigarette smoke also interacts with the lung through the airway epithelium, resulting in cellular and molecular changes (44), including dysregulation of EV’s secretion and bioactivity (45). Consistent with previous studies (46), we found during cigarette smoke exposure pro-inflammatory cytokines, such as TNF-α and IL-1β, associated with EVs are released from airway epithelial cells. In addition, cigarette smoke suppressed the release of EV-associated IL-1Ra, an IL-1β specific receptor antagonist. In COPD, either overproduction of IL-1 and/or underproduction of IL-1Ra can trigger a pro-inflammatory phenotype (47). Our results support the previous findings that pro-inflammatory cargo in cigarette smoke-induced EVs released by airway epithelial cells may contribute to lung injuries mediated by cigarette smoke.

In basal conditions, AATD macrophages produce higher amounts of pro-inflammatory cytokines, as well as signs of ER stress, relative to normal macrophages, suggesting their pro-inflammatory phenotype and lower activation threshold. This led us to hypothesize that AATD macrophages may be influenced by cigarette smoke-induced EVs due to their lower basal activation state and dysregulated immune responses. We found that neither control nor cigarette smoke-induced EVs released by airway epithelial cells had an effect on cytokine production by normal macrophages, while AATD macrophages were found to produce greater amounts of GM-CSF and IL-8 in response to cigarette smoke-induced EVs. It is well established that IL-8, a major chemoattractant for neutrophils, has a pivotal role in acute exacerbations of COPD (48). GM-CSF also has been shown to bind to the GM-CSF receptor on macrophages, enhancing macrophage proliferation and activation of the NF-κB pathway as an autocrine effect (49). Therefore, NF-κB appears to play a pivotal role in inflammatory processes by upregulating transcription of pro-inflammatory cytokine genes in response to stimuli (4, 27, 50). Our results indicate activation of the NF-κB pathway in AATD macrophages exposed to cigarette smoke. This suggests NF-κB activation may play a role in the response of AATD macrophages to cigarette smoke-induced EVs whereas inhibition of the NF-κB pathway represses this exacerbated response of AATD macrophages. Our results also revealed that cigarette smoke-induced EVs are able to induce ER stress in normal macrophages, while no changes were observed in ER stress levels of AATD macrophages. One possible explanation is that the levels of ER stress in AATD macrophages is already high and can mask the ER effect of cigarette smoke-induced EVs in AATD macrophages. These data suggest that in addition to disturbing the cytokine production in macrophages, cigarette smoke activates AATD macrophages via an unappreciated mechanism. Our data suggest that cigarette smoke-induced EVs released by other cell populations within the lung of AATD individuals may influence the secretion of pro-inflammatory mediators from AATD macrophages that can target downstream inflammatory signaling cascades during lung inflammation.

We have also found elevated levels of secreted ZAAT polymers in the conditioned media from AATD macrophages in response to cigarette smoke-induced EVs. This suggests the accumulation of ZAAT, in addition to NF-κB activation, may overwhelm the proteasome, leading to secretion of polymers (20). This is potentially relevant to lung inflammation since ZAAT polymers act as neutrophil chemoattractants and can mediate neutrophil degranulation (33).

Airway epithelial cells have been shown to synthesize and secrete small amounts of AAT protein, which may contribute to the pathogenesis of COPD in AATD individuals (51). A limitation in our study is that we were unable to obtain airway epithelial cells with an AATD phenotype to compare their EVs to normal airway epithelial cell-derived EVs. This limitation is difficult to overcome, given the limited access to patient-derived cell populations for rare diseases, including AATD. In conclusion our study reveals that expression of ZAAT contributes to the pro-inflammatory phenotype of AATD macrophages. We have shown that AATD macrophages have higher basal production of pro-inflammatory cytokines and a lower activation threshold, resulting in a dysregulated immune response to low levels of stimulation. This pro-inflammatory phenotype triggers release of GM-CSF and IL-8 from AATD macrophages in response to cigarette smoke-induced EVs released from airway epithelial cells. Release of these cytokines may play a role in neutrophil recruitment to the lung and inflammation in AATD individuals. Cigarette smoke-induced EVs also induce the release of ZAAT polymers from AATD macrophages, which have been shown to act as chemotactic signals for neutrophil recruitment to the lung. In this regard, the mechanism presented here may help us to better understand the multifaceted effect of AATD on lung tissue homeostasis. This mechanism can be critical to develop improved therapies for lung inflammation associated with AATD. These findings may translate into identifying a novel strategy to control enhanced AATD macrophage response to cigarette smoke in AATD individuals with lung inflammation.

## Abbreviations

AAT: alpha-1 antitrypsin
AATD: alpha-1 antitrypsin deficiency
COPD: chronic obstructive pulmonary disease
ER: endoplasmic reticulum
EV: extracellular vesicle
GM-CSF: granulocyte macrophage-colony stimulating factor
M-CSF: macrophage-colony stimulating factor
MDM: monocyte-derived macrophage
NE: neutrophil elastase
PBMC: peripheral blood mononuclear cell.

## Acknowledgments

We would like to thank the members of our laboratories for their contribution to various aspects of our research. The ultrastructural studies were conducted at the University of Florida College of Medicine Electron Microscopy Core Facility under the kindly supervision of Dr. Jill Verlander with the assistance of Dr. Sharon Matthews.

## Ethics approval and consent to participate

All participants gave informed, written consent prior to enrolling. The study was approved by the University of Florida institutional IRB (IRB # # 2015-01051).

## Consent for publication

Not Applicable.

## Availability of data and materials

The samples, datasets and analysis of this study are available from the corresponding author on reasonable request.

## Authors’ contributions

N.K. – study concept and design, acquisition of data, analysis and interpretation of data, statistical analysis, drafting of manuscript, study supervision. R.O. – acquisition of data, technical support. B.M. – analysis and interpretation of data, critical revision of manuscript for important intellectual content, study supervision. J.E.L. – acquisition of data, analysis, and interpretation of data. X.Q. – technical support. J.R.W. – acquisition of data. L.S.H. – analysis and interpretation of data, critical revision of manuscript for important intellectual content, drafting of manuscript. J.L. – acquisition and interpretation of data. G.W. – technical support. S.E. – drafting of manuscript, preparing the figures. M.B. – study concept and design, analysis and interpretation of the data, critical revision of manuscript for important intellectual content, obtained funding, and study supervision. All authors read and approved the final manuscript.

## References

1. Khodayari N, Oshins R, Holliday LS, Clark V, Xiao Q, Marek G, et al. Alpha-1 antitrypsin deficient individuals have circulating extracellular vesicles with profibrogenic cargo. Cell Commun Signal. 2020;18(1):140.

2. Krotova K, Marek GW, Wang RL, Aslanidi G, Hoffman BE, Khodayari N, et al. Alpha-1 Antitrypsin-Deficient Macrophages Have Increased Matriptase-Mediated Proteolytic Activity. Am J Respir Cell Mol Biol. 2017;57(2):238–47.

3. Di Stefano A, Caramori G, Oates T, Capelli A, Lusuardi M, Gnemmi I, et al. Increased expression of nuclear factor-kappaB in bronchial biopsies from smokers and patients with COPD. Eur Respir J. 2002;20(3):556–63.

4. Schuliga M. NF-kappaB Signaling in Chronic Inflammatory Airway Disease. Biomolecules. 2015;5(3):1266–83.

5. Lee J, Lu Y, Oshins R, West J, Moneypenny CG, Han K, et al. Alpha 1 Antitrypsin-Deficient Macrophages Have Impaired Efferocytosis of Apoptotic Neutrophils. Front Immunol. 2020;11:574410.

6. Van’t Wout EF, van Schadewijk A, Lomas DA, Stolk J, Marciniak SJ, Hiemstra PS. Function of monocytes and monocyte-derived macrophages in alpha1-antitrypsin deficiency. Eur Respir J. 2015;45(2):365–76.

7. Massari S, Nannetti G, Goracci L, Sancineto L, Muratore G, Sabatini S, et al. Structural investigation of cycloheptathiophene-3-carboxamide derivatives targeting influenza virus polymerase assembly. J Med Chem. 2013;56(24):10118–31.

8. Li G, Liu Y, Xie C, Zhou Q, Chen X. Characteristics of expedited programmes for cancer drug approval in China. Nat Rev Drug Discov. 2021;20(6):416.

9. Belchamber KBR, Donnelly LE. Macrophage Dysfunction in Respiratory Disease. Results Probl Cell Differ. 2017;62:299–313.

10. Bergin DA, Reeves EP, Meleady P, Henry M, McElvaney OJ, Carroll TP, et al. alpha-1 Antitrypsin regulates human neutrophil chemotaxis induced by soluble immune complexes and IL-8. J Clin Invest. 2010;120(12):4236–50.

11. Belchamber KBR, Walker EM, Stockley RA, Sapey E. Monocytes and Macrophages in Alpha-1 Antitrypsin Deficiency. Int J Chron Obstruct Pulmon Dis. 2020;15:3183–92.

12. Mulgrew AT, Taggart CC, Lawless MW, Greene CM, Brantly ML, O’Neill SJ, et al. Z alpha1-antitrypsin polymerizes in the lung and acts as a neutrophil chemoattractant. Chest. 2004;125(5):1952–7.

13. Corsello T, Kudlicki AS, Garofalo RP, Casola A. Cigarette Smoke Condensate Exposure Changes RNA Content of Extracellular Vesicles Released from Small Airway Epithelial Cells. Cells. 2019;8(12).

14. Bazzan E, Radu CM, Tine M, Neri T, Biondini D, Semenzato U, et al. Microvesicles in bronchoalveolar lavage as a potential biomarker of COPD. Am J Physiol Lung Cell Mol Physiol. 2021;320(2):L241–L5.

15. Wang N, Wang Q, Du T, Gabriel ANA, Wang X, Sun L, et al. The Potential Roles of Exosomes in Chronic Obstructive Pulmonary Disease. Front Med (Lausanne). 2020;7:618506.

16. Song Q, Chen P, Liu XM. The role of cigarette smoke-induced pulmonary vascular endothelial cell apoptosis in COPD. Respir Res. 2021;22(1):39.

17. Carroll TP, Greene CM, O’Connor CA, Nolan AM, O’Neill SJ, McElvaney NG. Evidence for unfolded protein response activation in monocytes from individuals with alpha-1 antitrypsin deficiency. J Immunol. 2010;184(8):4538–46.

18. Hu B, Liu J, Wu Z, Liu T, Ullenbruch MR, Ding L, et al. Reemergence of hedgehog mediates epithelial-mesenchymal crosstalk in pulmonary fibrosis. Am J Respir Cell Mol Biol. 2015;52(4):418–28.

19. Nasreen N, Khodayari N, Sriram PS, Patel J, Mohammed KA. Tobacco smoke induces epithelial barrier dysfunction via receptor EphA2 signaling. Am J Physiol Cell Physiol. 2014;306(12):C1154–66.

20. Khodayari N, Oshins R, Alli AA, Tuna KM, Holliday LS, Krotova K, et al. Modulation of calreticulin expression reveals a novel exosome-mediated mechanism of Z variant alpha1-antitrypsin disposal. J Biol Chem. 2019;294(16):6240–52.

21. Khodayari N, Wang RL, Marek G, Krotova K, Kirst M, Liu C, et al. SVIP regulates Z variant alpha-1 antitrypsin retro-translocation by inhibiting ubiquitin ligase gp78. PLoS One. 2017;12(3):e0172983.

22. Yang DC, Chen CH. Cigarette Smoking-Mediated Macrophage Reprogramming: Mechanistic Insights and Therapeutic Implications. J Nat Sci. 2018;4(11).

23. Ito H, Yamashita Y, Tanaka T, Takaki M, Le MN, Yoshida LM, et al. Cigarette smoke induces endoplasmic reticulum stress and suppresses efferocytosis through the activation of RhoA. Sci Rep. 2020;10(1):12620.

24. Elliott CL, Allport VC, Loudon JA, Wu GD, Bennett PR. Nuclear factor-kappa B is essential for up-regulation of interleukin-8 expression in human amnion and cervical epithelial cells. Mol Hum Reprod. 2001;7(8):787–90.

25. Schreck R, Baeuerle PA. NF-kappa B as inducible transcriptional activator of the granulocyte-macrophage colony-stimulating factor gene. Mol Cell Biol. 1990;10(3):1281–6.

26. Wang X, Lennard Richard M, Li P, Henry B, Schutt S, Yu XZ, et al. Expression of GM-CSF Is Regulated by Fli-1 Transcription Factor, a Potential Drug Target. J Immunol. 2021;206(1):59–66.

27. Joshi-Barve S, Barve SS, Butt W, Klein J, McClain CJ. Inhibition of proteasome function leads to NF-kappaB-independent IL-8 expression in human hepatocytes. Hepatology. 2003;38(5):1178–87.

28. Strzelak A, Ratajczak A, Adamiec A, Feleszko W. Tobacco Smoke Induces and Alters Immune Responses in the Lung Triggering Inflammation, Allergy, Asthma and Other Lung Diseases: A Mechanistic Review. Int J Environ Res Public Health. 2018;15(5).

29. Alam S, Li Z, Janciauskiene S, Mahadeva R. Oxidation of Z alpha1-antitrypsin by cigarette smoke induces polymerization: a novel mechanism of early-onset emphysema. Am J Respir Cell Mol Biol. 2011;45(2):261–9.

30. Mahadeva R, Atkinson C, Li Z, Stewart S, Janciauskiene S, Kelley DG, et al. Polymers of Z alpha1-antitrypsin co-localize with neutrophils in emphysematous alveoli and are chemotactic in vivo. Am J Pathol. 2005;166(2):377–86.

31. Wang L, Chen Q, Yu Q, Xiao J, Zhao H. Cigarette smoke extract-treated airway epithelial cells-derived exosomes promote M1 macrophage polarization in chronic obstructive pulmonary disease. Int Immunopharmacol. 2021;96:107700.

32. Ogger PP, Byrne AJ. Macrophage metabolic reprogramming during chronic lung disease. Mucosal Immunol. 2021;14(2):282–95.

33. Parmar JS, Mahadeva R, Reed BJ, Farahi N, Cadwallader KA, Keogan MT, et al. Polymers of alpha(1)-antitrypsin are chemotactic for human neutrophils: a new paradigm for the pathogenesis of emphysema. Am J Respir Cell Mol Biol. 2002;26(6):723–30.

34. Sukhorukov VN, Khotina VA, Bagheri Ekta M, Ivanova EA, Sobenin IA, Orekhov AN. Endoplasmic Reticulum Stress in Macrophages: The Vicious Circle of Lipid Accumulation and Pro-Inflammatory Response. Biomedicines. 2020;8(7).

35. Barnes PJ. The cytokine network in chronic obstructive pulmonary disease. Am J Respir Cell Mol Biol. 2009;41(6):631–8.

36. da Silva CO, Gicquel T, Daniel Y, Bartholo T, Vene E, Loyer P, et al. Alteration of immunophenotype of human macrophages and monocytes after exposure to cigarette smoke. Sci Rep. 2020;10(1):12796.

37. Grumelli S, Corry DB, Song LZ, Song L, Green L, Huh J, et al. An immune basis for lung parenchymal destruction in chronic obstructive pulmonary disease and emphysema. PLoS Med. 2004;1(1):e8.

38. Devereux G. ABC of chronic obstructive pulmonary disease. Definition, epidemiology, and risk factors. BMJ. 2006;332(7550):1142–4.

39. Jorgensen E, Stinson A, Shan L, Yang J, Gietl D, Albino AP. Cigarette smoke induces endoplasmic reticulum stress and the unfolded protein response in normal and malignant human lung cells. BMC Cancer. 2008;8:229.

40. Reverendo M, Mendes A, Arguello RJ, Gatti E, Pierre P. At the crossway of ER-stress and proinflammatory responses. FEBS J. 2019;286(2):297–310.

41. Bazzan E, Tine M, Biondini D, Benetti R, Baraldo S, Turato G, et al. alpha1-Antitrypsin Polymerizes in Alveolar Macrophages of Smokers With and Without alpha1-Antitrypsin Deficiency. Chest. 2018;154(3):607–16.

42. Gaschler GJ, Zavitz CC, Bauer CM, Skrtic M, Lindahl M, Robbins CS, et al. Cigarette smoke exposure attenuates cytokine production by mouse alveolar macrophages. Am J Respir Cell Mol Biol. 2008;38(2):218–26.

43. Alam S, Li Z, Atkinson C, Jonigk D, Janciauskiene S, Mahadeva R. Z alpha1-antitrypsin confers a proinflammatory phenotype that contributes to chronic obstructive pulmonary disease. Am J Respir Crit Care Med. 2014;189(8):909–31.

44. Goldfarbmuren KC, Jackson ND, Sajuthi SP, Dyjack N, Li KS, Rios CL, et al. Dissecting the cellular specificity of smoking effects and reconstructing lineages in the human airway epithelium. Nat Commun. 2020;11(1):2485.

45. Benedikter BJ, Volgers C, van Eijck PH, Wouters EFM, Savelkoul PHM, Reynaert NL, et al. Cigarette smoke extract induced exosome release is mediated by depletion of exofacial thiols and can be inhibited by thiol-antioxidants. Free Radic Biol Med. 2017;108:334–44.

46. Mohan A, Agarwal S, Clauss M, Britt NS, Dhillon NK. Extracellular vesicles: novel communicators in lung diseases. Respir Res. 2020;21(1):175.

47. Suwara MI, Green NJ, Borthwick LA, Mann J, Mayer-Barber KD, Barron L, et al. IL-1alpha released from damaged epithelial cells is sufficient and essential to trigger inflammatory responses in human lung fibroblasts. Mucosal Immunol. 2014;7(3):684–93.

48. Gilowska I. [CXCL8 (interleukin 8)--the key inflammatory mediator in chronic obstructive pulmonary disease?]. Postepy Hig Med Dosw (Online). 2014;68:842–50.

49. Martinez FO, Gordon S. The M1 and M2 paradigm of macrophage activation: time for reassessment. F1000Prime Rep. 2014;6:13.

50. Caramori G, Romagnoli M, Casolari P, Bellettato C, Casoni G, Boschetto P, et al. Nuclear localisation of p65 in sputum macrophages but not in sputum neutrophils during COPD exacerbations. Thorax. 2003;58(4):348–51.

51. Pini L, Tiberio L, Venkatesan N, Bezzi M, Corda L, Luisetti M, et al. The role of bronchial epithelial cells in the pathogenesis of COPD in Z-alpha-1 antitrypsin deficiency. Respir Res. 2014;15:112.

