## Supplementary figures and images for "Cigarette smoke exposed airway epithelial cell-derived EVs promote pro-inflammatory macrophage activation in alpha-1 antitrypsin deficiency"

### Supplementary Figure 1

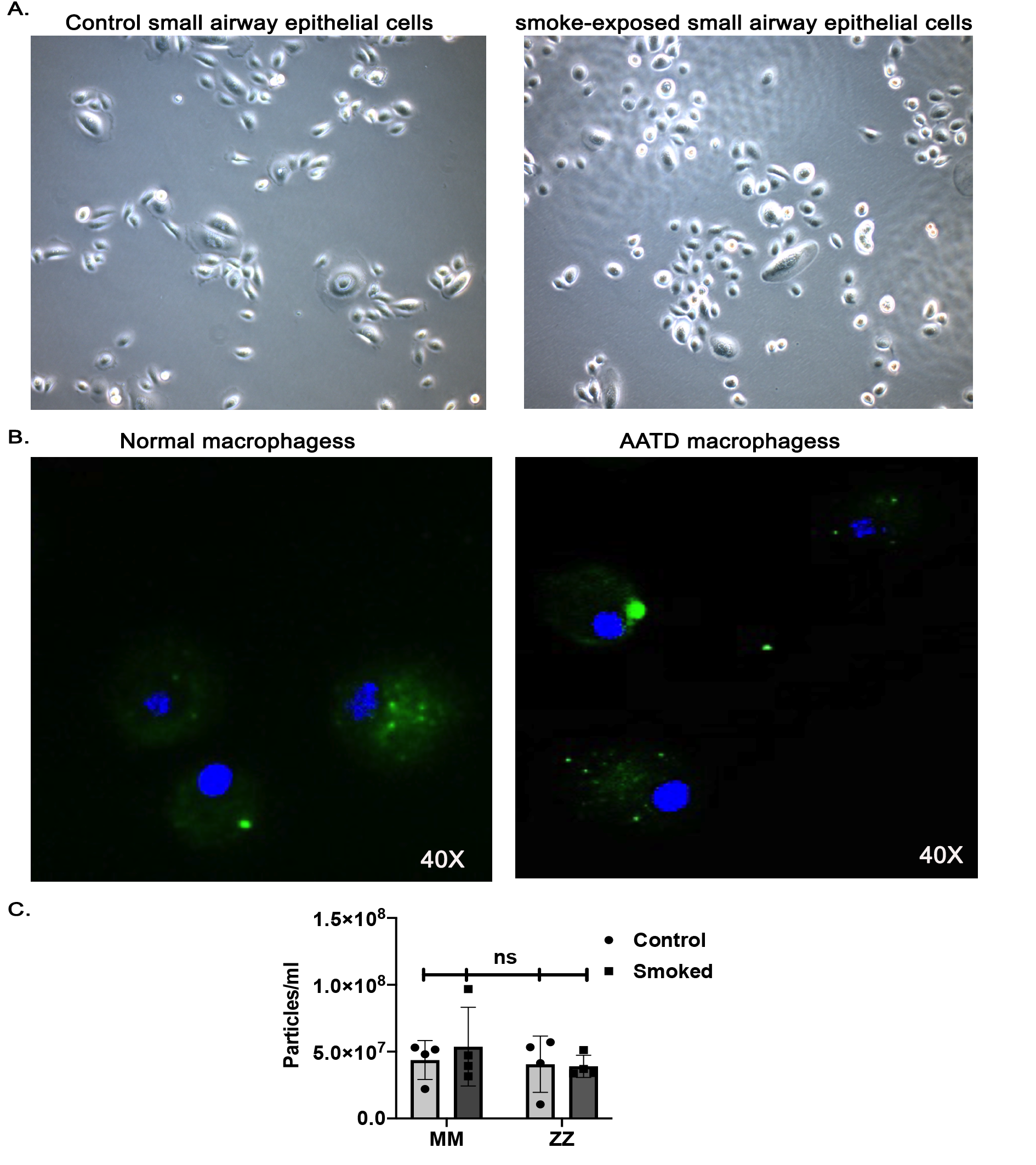

### Supplementary Figure 2

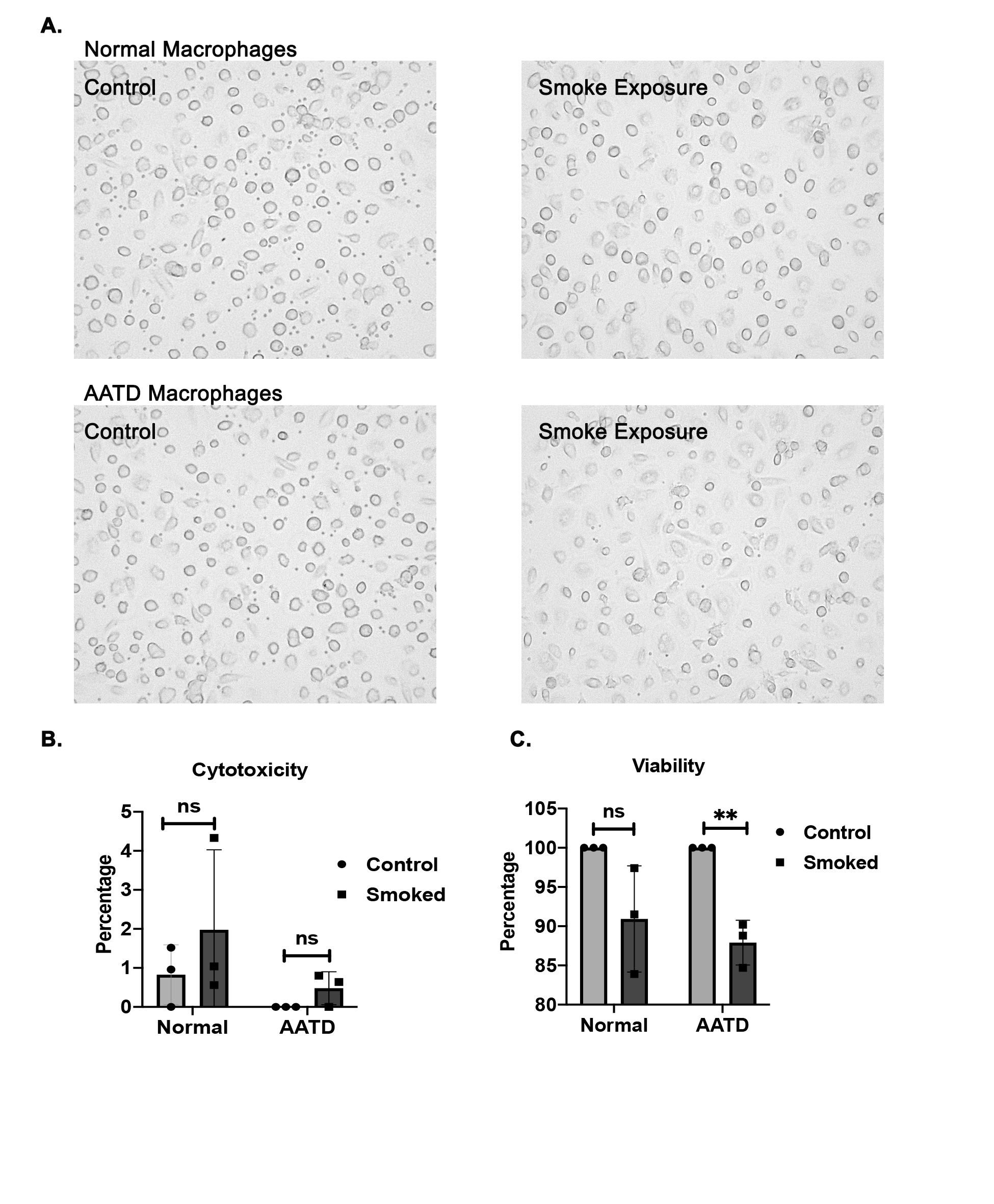

### Supplementary Figure 3

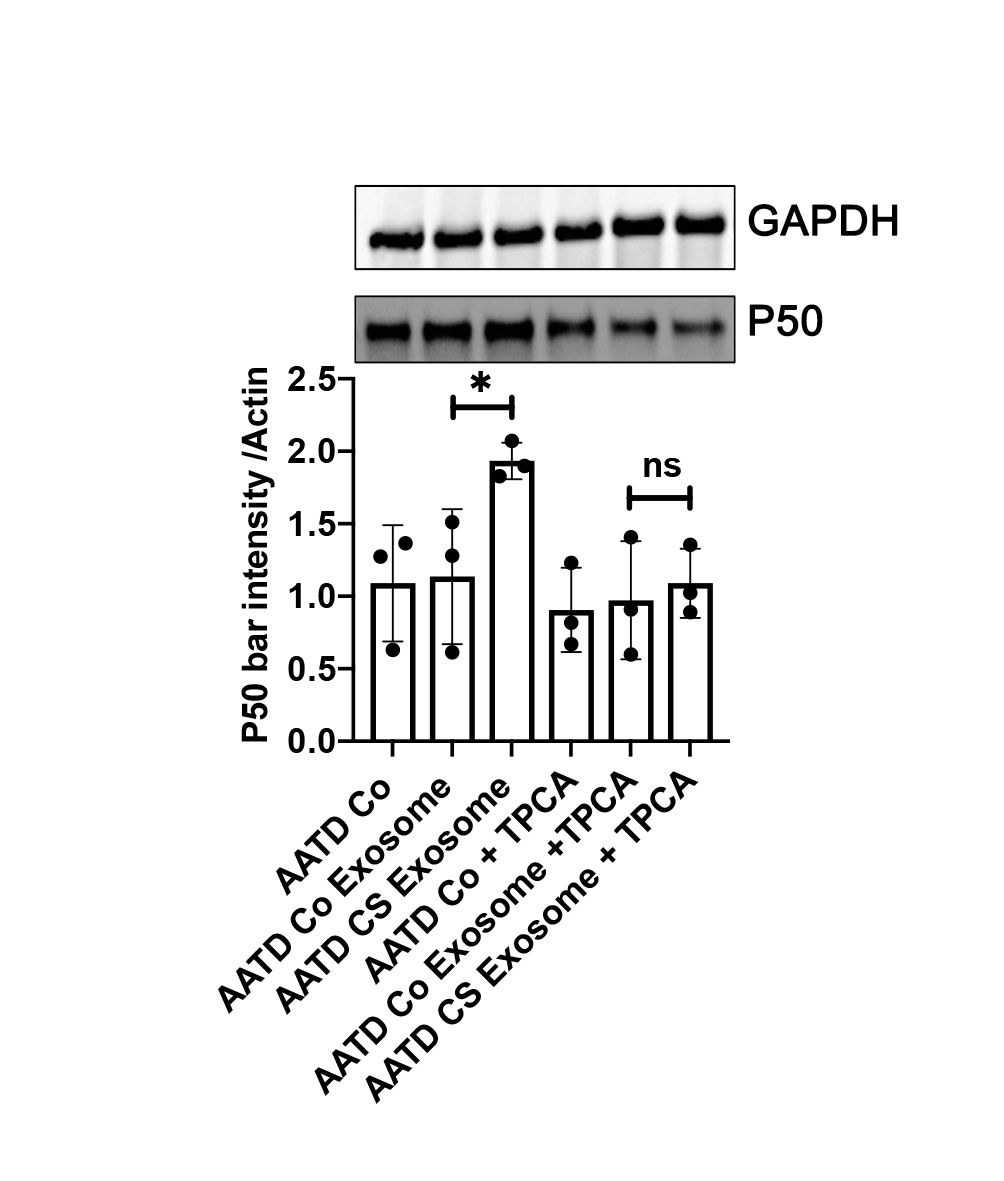
