## Supplemental Figure Legends for "Cigarette smoke exposed airway epithelial cell-derived EVs promote pro-inflammatory macrophage activation in alpha-1 antitrypsin deficiency"

Online Data Supplement

Supplementary Figure 1. (A) The morphology of small airway epithelial cells before and after smoke was monitored using light microscopy. (B) DIO labeled (green) exosomes were incubated with normal and AATD macrophages for 1 hour. The immunofluorescence pictures of trypsin treated cells shows green labeled exosomes inside the normal and AATD macrophages. (C) The concentration of EVs release by control and smoked normal (MM) and AATD (ZZ) macrophages as determined by Nano Track Analysis.

Supplementary Figure 2. (A) The morphology of normal and AATD macrophages before and after smoke was monitored using light microscopy. (B) LDH assay indicating the percentage of cytotoxicity and (C) MTT assay indicating viability of normal and AATD macrophages have been presented.

Supplementary Figure 3. The representative blot of protein levels of P50 in AATD macrophages with/without NF-κB inhibitor (TPCA) accompanied with normalized bar intensities to actin (n=3).
